# WHO WINS? COMPARISON BETWEEN EMERGENT MACROPHYTES AND PONTEDERIA CRASSIPES (WATER HYACINTH) AS NATURE-BASED SOLUTIONS FOR NUTRIENT PHYTOREMEDIATION AND CYANOBACTERIA MITIGATION

**DOI:** 10.1101/2024.07.16.603719

**Authors:** Cesar Macedo Lima Filho, Allan Amorim Santos, Daniel Vinícius Neves de Lima, Luan de Oliveira Silva, Ana Beatriz Furlanetto Pacheco, Sandra M. F. O. Azevedo

**Affiliations:** Laboratory of Ecophysiology and Toxicology of Cyanobacteria - Carlos Chagas Filho Institute of Biophysics, Federal University of Rio de Janeiro. Avenida Carlos Chagas Filho, 373, CCS-Bloco G, Ilha do Fundão, CEP: 21941-902, Rio de Janeiro, Brazil; Laboratory of Biological Physics - Carlos Chagas Filho Institute of Biophysics, Federal University of Rio de Janeiro. Avenida Carlos Chagas Filho, 373, CCS-Bloco G, Ilha do Fundão, CEP: 21941-902, Rio de Janeiro, Brazil

**Keywords:** Phytoremediation, emergent macrophyte, mesocosm, cyanobacteria, eutrophication

## Abstract

Emergent macrophytes are commonly used to control eutrophication in Constructed Floating Wetlands (CFW) systems. However, tropical emergent macrophyte species, particularly from South America, are underrepresented in these applications. This study aims to investigate the potential of five emergent macrophytes, *Alternanthera philoxeroides, Ludwigia leptocarpa, Polygonum ferrugineum, Typha domingensis,* and *Urochloa mutica* in controlling eutrophication and inhibiting cyanobacterial growth, compared to the floating macrophyte *Pontederia crassipes,* the most studied macrophyte for this purpose. Phytoremediation experiments were performed ex-situ in mesocosms (50 L) with high soluble reactive phosphorus (SRP) concentration (400 µg L^−1^). Allelopathic effects of the macrophytes root exudates on cyanobacteria were tested in vitro using a strain of *Microcystis aeruginosa*. While *P. crassipes* reduced SRP concentration by 86% in 14 days, the tested macrophytes reduced SRP concentration by > 94%, except for *T. domingensis,* which showed a reduction of 64%. NH_4_^+^ and NO_3_^−^ removal in 14 days were 96% and 62%, respectively, for *P. crassipes*. These values corresponded to 96% and 63% for the five tested macrophytes. Root exudates of *A. philoxeroides, L. leptocarpa,* and *P. ferrugineum* caused inhibition of *M. aeruginosa* growth with no detection of Chl-*a* after 7 days. Thus, three emergent macrophytes were more efficient in removing nutrients than *P. crassipes*. and showed allelopathic potential against cyanobacteria, indicating that using local emergent macrophytes in CFW systems can be a valuable and sustainable tool to mitigate eutrophication and its consequences in aquatic environments.

**Graphical abstract:** 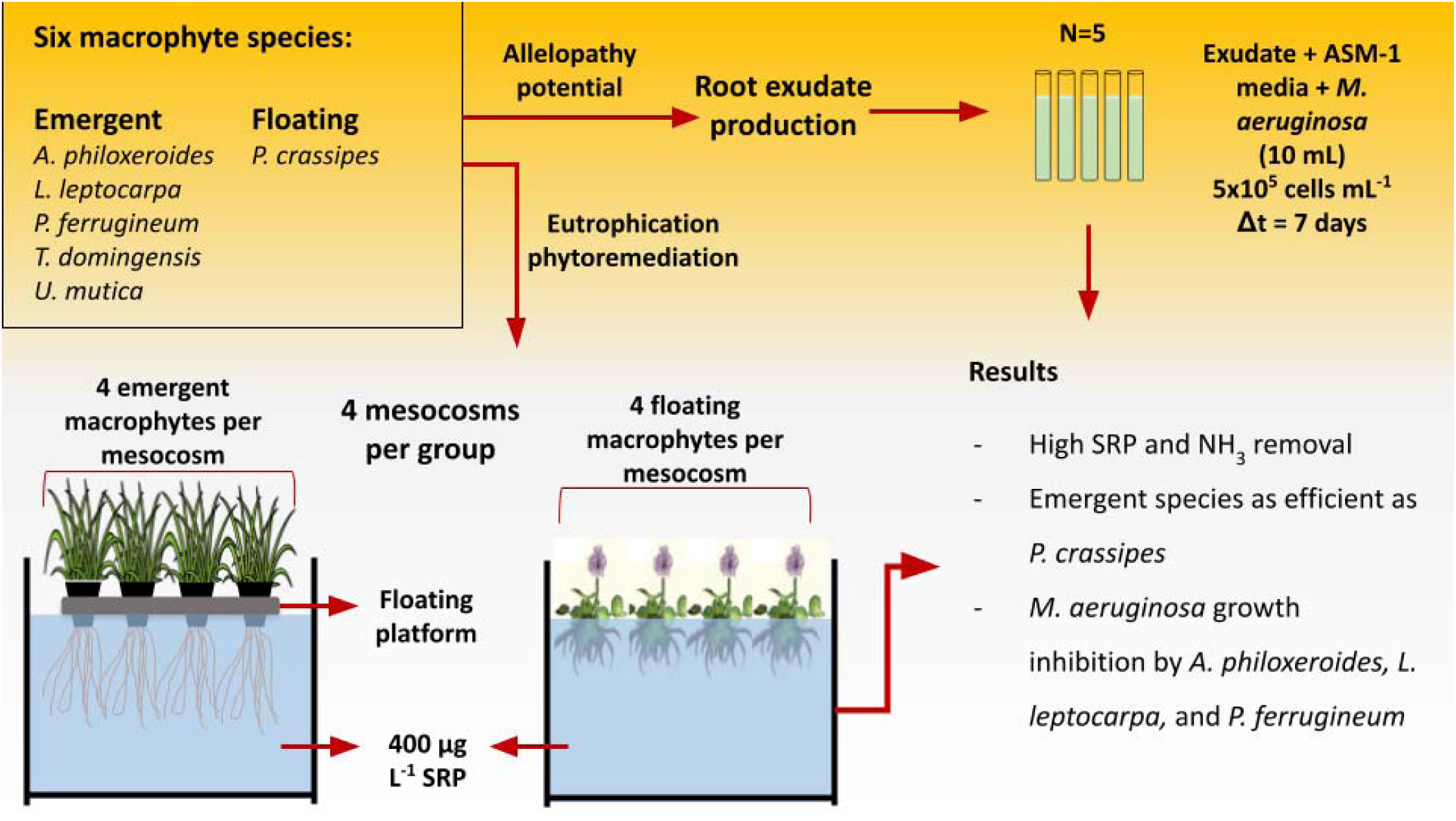

**Highlights:**

- Emergent macrophytes were similarly efficient to *P. crassipes* in nutrient removal
- *Ludwigia leptocarpa* was the most efficient in SRP removal in 7 days
- Allelopathy on cyanobacterium *M. aeruginosa* by *A. philoxeroides, L. leptocarpa*, and *P. ferrugineum* root exudates
- Adapting emergent macrophytes in CFW structures can be helpful in in situ phytoremediation.

**Statement of Novelty:** This study lies in its comparative analysis of underrepresented tropical emergent macrophytes from South America against the widely studied floating macrophyte, *Pontederia crassipes* (water hyacinth). By focusing on these local emergent macrophytes, the research addresses a significant knowledge gap. It provides insights into their potential application in Constructed Floating Wetlands (CFWs) for nutrient removal and cyanobacteria control. Key findings demonstrate that the selected emergent macrophytes outperform *P. crassipes* in reducing soluble reactive phosphorus (SRP) concentrations and exhibit substantial nitrogen removal efficiencies. Furthermore, the allelopathic potential of *A. philoxeroides*, *L. leptocarpa*, and *P. ferrugineum* against the harmful cyanobacterium *Microcystis aeruginosa* showcases their dual functionality in mitigating eutrophication and inhibiting harmful algal blooms.

## 1. Introduction

The intensification of agricultural and urban runoff has contributed to eutrophication due to increased nutrient loadings enriching aquatic ecosystems with nitrogen and phosphorus (Smith and Schindler, 2009; Le Moal et al., 2019; Akinnawo, 2023). Once these nutrients are available, they can be used by phytoplankton organisms and/or macrophytes, contributing to their excessive growth, unbalancing the trophic web and decreasing water quality (Paerl et al., 2011).

Among phytoplankton groups, cyanobacteria frequently cause blooms in eutrophic environments, and some genera can produce toxic metabolites that affect animals and humans, representing an additional risk (Polyak and Polyak, 2022). The cosmopolitan genus *Microcystis* is one of the most studied among freshwater bloom-forming cyanobacteria. It presents cells organized in colonies that protect from stress and grazing (Gobler et al., 2016). Because they are not diazotrophic, the availability of dissolved N is a limiting nutritional factor for their growth and is related to the formation of blooms. The availability of P is also a limiting factor, although *Microcystis* has adaptive strategies that allow growth in limiting concentrations of dissolved P (Gobler et al., 2016). Therefore, controlling the availability of N and P in aquatic systems is a basic strategy to minimize the occurrence of blooms of this genus as well as other cyanobacteria, contributing to improving the water quality (Hamilton et al., 2016; Paerl et al., 2020).

Several alternatives have been raised for controlling external and internal nutrient loading in aquatic environments and mitigating cyanobacterial blooms. For example, the use of lanthanum-containing materials to bind and sediment phosphate (Zeller & Alperin, 2021) or the application of artificial mixing of the water column to eliminate stratification, promote oxygenation, and reduce nutrient concentrations (Chen et al., 2024). However, the application of chemical and physical methods is challenging due to operational costs, the need for stakeholder involvement and scale limitations (Shortle et al., 2020). Biomanipulation methods can be used to control eutrophication and reduce cyanobacterial blooms, such as introducing planktivorous organisms or macrophytes in aquatic systems. These constitute nature-based solutions (NBS) that can increase biodiversity, restabilize aquatic environments and recover ecosystem services (Triest et al., 2016; Eggermont et al., 2015; Boelee et al., 2017)

Phytoremediation, a bioprocess that uses plants to remove pollutants from soil or water, is a sustainable NBS tool for eutrophication control that takes advantage of a physiological process of the organism (Kumar et al., 2018). For aquatic plants (macrophytes) phytoremediation can be combined with allelopathy, potentially controlling cyanobacterial growth (Mohamed, 2017). Floating macrophytes such as *Pontederia crassipes* (water hyacinth) and *Pistia stratiotes* (water lettuce) are commonly used in phytoremediation because they present high nutrient removal capacity and can thrive in polluted environments (Henares and Camargo, 2014; Qin et al., 2016b; Mishra and Maiti, 2017; Osti et al., 2020; Auchterlonie et al., 2021). These macrophytes also exert allelopathic effects on cyanobacteria (Wu et al., 2015, 2019; Lourenção et al., 2021; De Lima et al., 2023). Nonetheless, their fast and potentially uncontrolled spread over the water surface is an environmental, social, and economic problem (Datta et al., 2021). Despite the advantages of macrophyte introduction to control eutrophication, it is not recommended to introduce non-native species in phytoremediation efforts due to the risk of invasiveness (Patel, 2012; Djihouessi et al., 2023).

Constructed Floating Wetlands (CFW) are systems that use emergent aquatic macrophytes in floating platforms for phytoremediation, mimicking the behaviour of floating macrophytes. In these systems, the emergent macrophyte roots are exposed to absorb the nutrients from the water column. CFW systems are inspired by natural floating wetlands, buoyant islands of organic matter where emergent macrophytes can grow, shifting from a terrestrial to a floating behaviour (Sasser et al., 1991). The macrophyte roots have a central role in the system’s function, and their association with microbes optimizes several biochemical processes and nutrient cycling (Shahid et al., 2018). The efficiency of CFW in nutrient removal and phytoplankton growth control is well documented but mainly for Eurasian macrophyte species from temperate climates (Geng et al., 2017; West et al., 2017; Schwammberger et al., 2019; Vymazal, 2022). This points to the potential to explore a greater diversity of plants from other geographic regions. Considering that South America hosts high biodiversity and faces serious eutrophication problems, investigation of the phytoremediation potential of native species can contribute to expanding the use of CFW (Rolon and Maltchik, 2006; Ferreira et al., 2011) as well as to explore allelopathic effects on cyanobacteria. Ideally, CFW should use local macrophytes to avoid introducing exotic species (Lucke et al., 2019) and its development with native plants constitutes an interesting NBS to control eutrophication and cyanobacterial blooms.

Considering these previous observations, we chose five emergent macrophytes species to be tested in this study: *Alternanthera philoxeroides* (Mart.) Griseb., *Ludwigia leptocarpa* (Nutt.) H. Hara, *Polygonum ferrugineum* Wedd., *Typha domingensis* Pers. and *Urochloa mutica* (Forssk.) T.Q.Nguyen. These species have different ecophysiological traits that allow them to colonize the aquatic environment, being easily propagated. They are distributed mainly in South America and Central America, although some can also be found in North America. Regarding Liliopsida monocotyledonous, *T. domingensis* thrives in flooded areas occupying degraded sites with dense vegetation through its rhizomes (Cruz et al., 2019) while *U. mutica* is a grass native to Africa, although naturalized in Brazil, which spreads over the substrate and adapts to both flooded and dry conditions (Williams and Baruch, 2000; Martins et al., 2008; de Moura-Júnior et al., 2013). Regarding Magnoliopsida dicotyledonous, *A. philoxeroides* is a perennial species with stolons and rhizomes, featuring floating stems due to stem aerenchyma, thriving in aquatic margins, flooded areas, and dry land, reproducing vegetatively (Griseb, 2016) as well as *P. ferrugineum* that creates floating branches when flooded (de Assis Murillo et al., 2019; dos Santos Machado et al., 2021), whereas *L. leptocarpa* inhabiting riverbanks, lakes, and flooded areas (De Sousa et al., 2019).

This study aimed to investigate the potential for nutrient removal by South American emergent macrophyte species in CFW systems and the allelopathic activity of macrophyte root exudates on a cyanobacterial species (*Microcystis aeruginosa*). The emergent macrophytes were compared to the commonly studied floating macrophyte *Pontederia crassipes* (Mart.) Solms. We hypothesized that the emergent macrophytes could be adapted to CFWs mimicking floating macrophytes and achieve similar or superior results in nutrient phytoremediation and cyanobacterial suppression.

## 2. Material and methods

### 2.1 Macrophyte and cyanobacteria cultivation

The six species of macrophytes were collected in natural environments. *A. philoxeroides*, *L. leptocarpa*, *P. ferrugineum*, and *P. crassipes* were collected in the Barra do Braúna reservoir (MG, Brazil) (21°26’50.83” S; 42°24’26.07” O) in 2019. *T. domingensis* and *U. mutica* were collected in the Santo Antônio reservoir (RJ, Brazil) (22° 8’59.66” S; 42°21’0.89” O) in 2020. Their taxonomical classification and distribution in the American continent are detailed in Table 1.

**Table 1:**
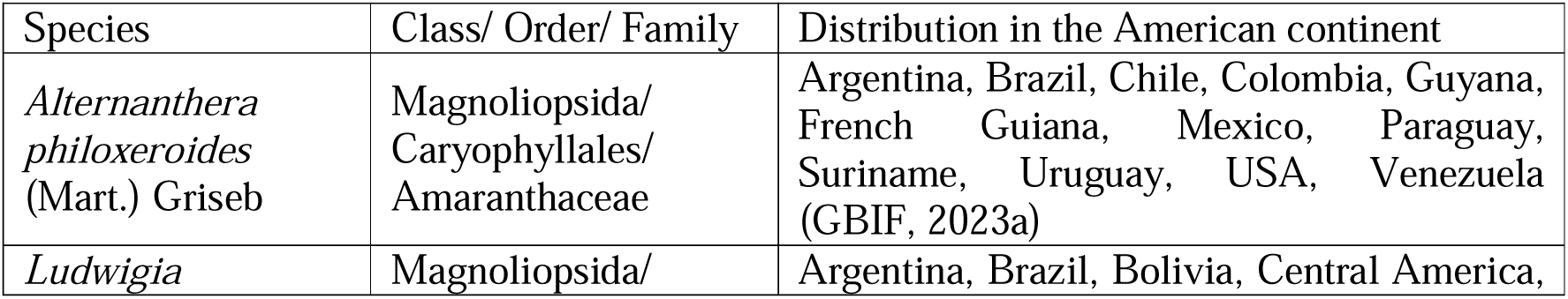
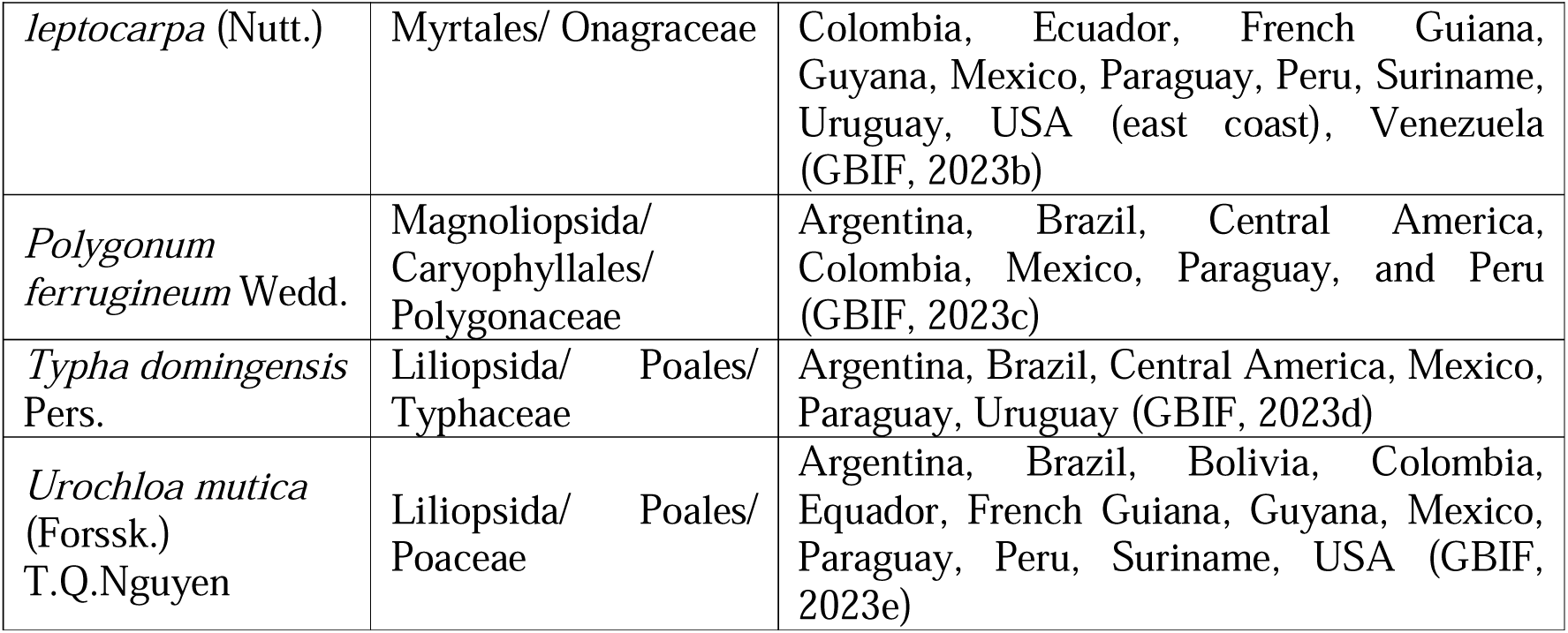
Taxonomical classification and distribution of the emergent macrophytes species chosen for this study.

Parent crops were maintained in a greenhouse located at the campus of the Federal University of Rio de Janeiro. *A. philoxeroides*, *P. ferrugineum*, and *U. mutica* were propagated by cuttings, *P. crassipes* and *T. domingensis* were propagated by separation of individuals from rhizomes and *L. leptocarpa* was propagated by seeds. The generated individuals were maintained in bags filled with water-saturated soil. After growth, they were removed from the substrate, and the roots were washed with tap water. Their aerial parts were trimmed, and the plants were transferred to flowering pots from which the bottom was removed for the roots to pass freely. A polyurethane foam was used to support the macrophytes. The pots fit on a polystyrene floating platform and were acclimated in a tank with tap water for 20 days. *P. crassipes* individuals were also maintained in tanks with tap water.

The cyanobacterium *Microcystis aeruginosa* (strain LETC-MC-02) was maintained in the collection of the Laboratory of Ecophysiology and Toxicology of Cyanobacteria. This strain was isolated from an urban lagoon (Jacarepaguá Lagoon, RJ, Brazil) (22°59’5.10” S 43°24’41.01” O). Cultures were routinely maintained in ASM-1 media (Gorham et al., 1964) at 25 °C, 50 µmol photon m^2^ s^−1,^ and 12h/12h light and dark photoperiod.

### 2.2 Mesocosm experimental design

The mesocosm experiment was conducted in the same greenhouse where the parent crops were maintained (Figure 1). The mesocosms (50 L) were filled with tap water. The inorganic fertilizer NPK 15:5:10 (Forth, Brazil) was added as a nutrient source to increase the soluble reactive phosphorus (SRP) concentration to around 400 µg L^−1^, simulating a hypereutrophic condition. Four mesocosms (N=4) for each experimental group were used, with four macrophyte individuals per mesocosm. The macrophytes were chosen by their similar size, with both aerial parts and roots approximately 20 cm in length, resulting in mean fresh biomass of 220.4 ± 33.0 g (*P. crassipes*), 261.2 ± 22.8 g (*A. philoxeroides*), 292.0 ± 31.6 g (*L. leptocarpa*), 235.1 ± 55.3 g (*P. ferrugineum*), 469.7 ± 50.8 g (*T. domingensis*), and 402.7 ± 37.2 g (*U. mutica*).

**Figure 1:**
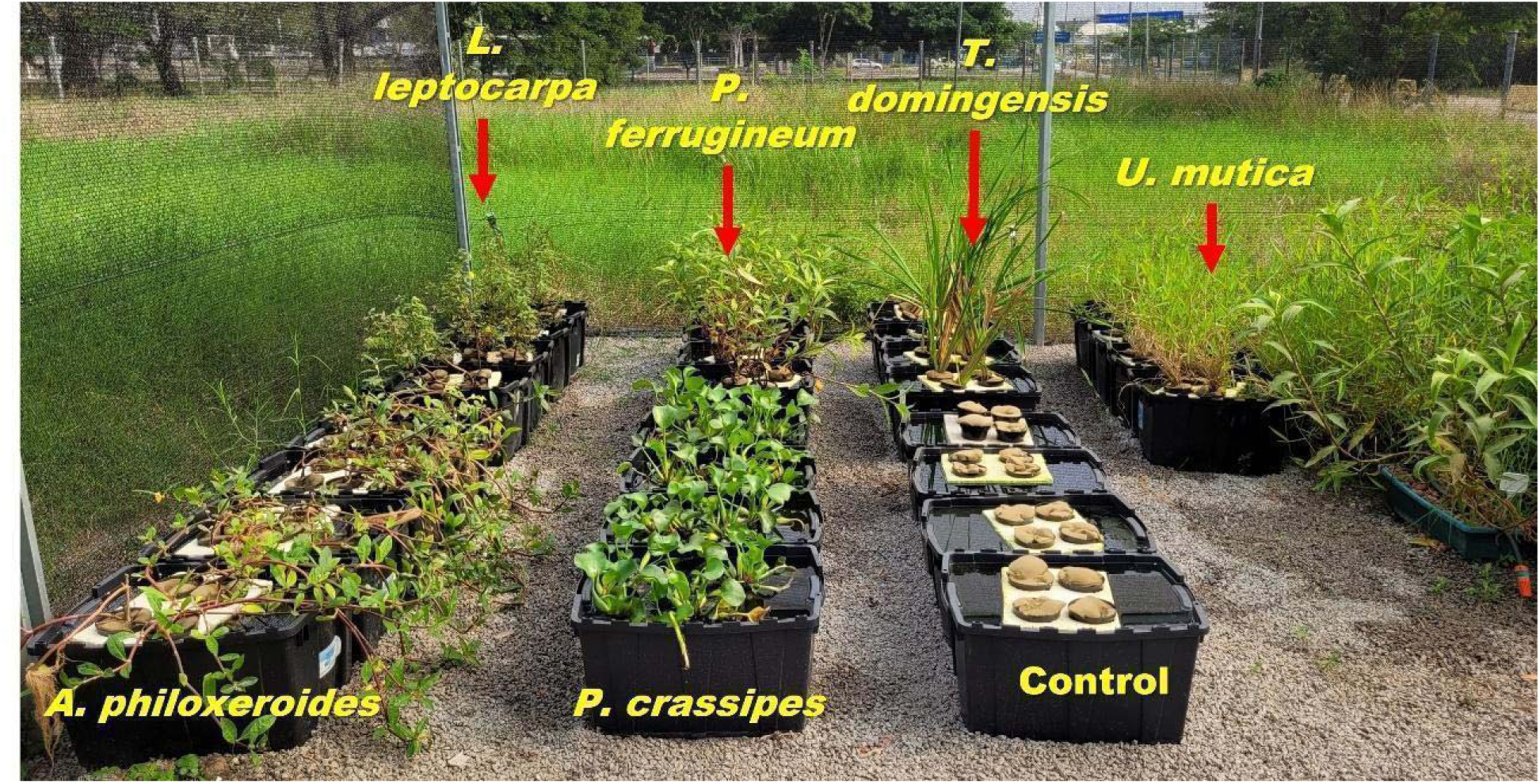
Cultivation of macrophytes in the greenhouse during the mesocosms experiment.

To build the CFWs, the macrophyte pots (obtained as described in section 2.1) were placed in a polystyrene floating platform (20 cm x 30 cm) with four holes to fit the pots. No platforms were used for *P. crassipes*, since it floats in water. A control group was set with flowering pots filled only with foam and placed on the platforms. The experiment was carried out between March and April 2021 and lasted for 14 days. Samples were taken at days 0 (start of the experiment), 7, and 14 (Figure 1).

### 2.3 Analysis of abiotic parameters in the water

Water samples were collected from mesocosms in plastic bottles and used to determine nutrient concentrations. The aliquots used to analyze dissolved nutrients (SRP, NH_4_^+^, and NO_3_^−^) were filtered on cellulose filters. The aliquots obtained to determine total phosphorus (TP) and total nitrogen (TN) concentrations were digested by adding 1 mL or 2.5 mL of 0.18 M potassium persulfate solution, respectively. SRP and TP concentrations were analyzed by the colorimetric method of Strickland and Parsons (1965), adding 1 mL of mixed solution to 10 ml of sample. The detection and quantification limits were 8 µg L^−1^ and 16 µg L^−1^, respectively. A standard curve was made by the serial dilution of a potassium phosphate solution. Volumes taken from the mesocosms for sampling or lost by evaporation were not replaced.

Nitrogen fractions were measured with the electrodes ISENH3181 (NH_4_^+^) and ISENO3181 (NO_3_^−^ and TN), both utilizing the multimeter probe Hatch HQ40D (Hach, USA), as recommended by the manufacturer (Standard Methods for the Examination of Water and Wastewater, 2018a, 2018b). The detection limits for NH_4_^+^ and NO_3_^−^ were 10 µg L^−1^ and 100 µg L^−1^, respectively.

Water conductivity, pH, dissolved oxygen (DO), and water temperature were measured by the multimeter probe Hatch HQ40D (Hach, USA) with the electrodes IntelliCAL CDC401 (conductivity), IntelliCAL PHC101 (pH), IntelliCAL LDO101 (DO). Air temperature was measured with an electronic thermometer.

### 2.4 In vitro allelopathy assay

Root exudates were prepared from living macrophytes (four individuals of each species) cultivated as described in section 2.1. The roots of each macrophyte were put into a glass flask (3 L) filled with Mili-Q water in a proportion of 100 g of biomass L^−1^, while the aerial part was left outside the flask. The top opening of the flask was covered with foam to help support the macrophytes, and the glass was covered with a black polyethylene layer to avoid light penetration and inhibit the growth of photosynthetic organisms. The macrophytes were kept outdoors for five days in this condition (mean temperature: 29 °C). The exudates produced were filtered in cellulose filters and frozen at −20 °C.

To test the allelopathic potential of root exudates against *M. aeruginosa*, the cyanobacterial culture medium ASM-1 (Gorham et al., 1964) was prepared using the root exudates instead of water to dilute the medium components. The exudates were thawed, filtered through 0.2 µm nylon membranes and added with ASM-1 salts, constituting a modified ASM-1 used in the treatment condition. By incorporating root exudates into the cyanobacterial culture medium, we intended to mimic the conditions found in freshwater systems, where macrophytes release compounds in the presence of cyanobacteria. In the control condition, cyanobacterial cultures were maintained in ASM-1. Cultures of *M. aeruginosa* (section 2.1) were diluted to constitute an inoculum of 5.0 x 10^5^ cells mL^−1^ in a final volume of 10 mL, kept in glass tubes maintained at 25 °C, 50 µmol photon m^2^ s^−1,^ and 12h/12h light and dark photoperiod. For each condition, 5 cultures were established.

At initial (t=0) and final (t=7 days) times, chlorophyll-*a* (Chl-*a*) concentrations and photosynthetic efficiency (yield) were measured using the fluorimeter Phyto-PAM (Walz, Germany), with a detection limit of 0.5 µg L^−1^. The effective quantum yield of photosynthetic energy conversion in photosystem II (PSII Yield) was determined considering the following equation (Genty et al., 1989):

Yield PSII = (Fm – F) / Fm,

F is the transient original fluorescence obtained before applying the saturation pulse, and Fm is the minimal fluorescence obtained when emitting a low-intensity measurement light.

### 2.5 Macrophyte and phytoplankton biomass

The fresh biomass of the macrophytes was measured by gravimetry at initial (t=0) and final (t=14 days) experimental days. Macrophytes were removed from the water, and the roots were allowed to drain for 10 minutes. The macrophyte growth ratio was calculated as Bf/Bi, where Bf was the final fresh biomass (after 14 days) and Bi was the fresh biomass at the initial time (t=0). The amount of P potentially assimilated by the macrophytes was estimated based on the difference between the initial concentration of dissolved P (SRP at T0) and the final concentration of dissolved P (SRP at T=14 days). The macrophyte biomass accumulation during this period was considered for this calculation as follows:

Ratio PO_4_^−3^: biomass = (Pi – Pf) / (Bf – Bi),

Where Pi is the initial SRP concentration, Pf is the final SRP concentration (14^th^ day), Bf is the final biomass (14^th^ day) and Bi is the initial biomass.

Phytoplankton growth was monitored in the mesocosm water during the experiment. It resulted from the water used to fill the mesocosms. The phytoplankton biomass was estimated by chlorophyll-*a* (Chl-*a*) concentration using Phyto-PAM (Walz, Germany). In this equipment, the Chl-*a* pigment is excited at four different wavelengths, which makes it possible to distinguish phytoplankton groups, considering each light-harvesting pigment antennae from Blue-Green Algae (Cyanophyta/Cyanobacteria), Green algae (Chlorophyta), and Brown Algae (Diatoms and/or Cryptophyta). The detection limit for Chl-*a* was 0.5 µg L^−1^. Additionally, PSII Yield was determined for each phytoplankton group, as described for *M. aeruginosa*.

### 2.6 Characterization of macrophyte root exudates by high-resolution mass spectrometry

The macrophyte root exudates (section 2.4) were lyophilized in 4 aliquots of 50 mL, each one originating from an individual macrophyte from each species. The material was suspended in water/acetonitrile 1:1 to a final concentration of 1 mg mL^−1^ and filtered through 0.22 µm cellulose membranes. Analyses were carried out on a Bruker SolariX XR, 7 Tesla Fourier Transform Ion Cyclotron Resonance Mass Spectrometry (FT-ICR-MS) (Brüker Daltonics Inc., Billerica, MA) equipped with an electrospray ionization (ESI) source and detailed information about the method is attached in Supplementary Material.

### 2.7 Statistical analysis

After basic assumptions such as normality and homoscedasticity were accepted, two-way ANOVA tests were applied to compare treatment and control conditions over time, considering both phytoremediation mesocosms and allelopathy experiments (p < 0.05). For single dependent variables, a one-way ANOVA was applied considering macrophyte biomass ratio (ordinary one-way ANOVA assuming equal variances) as well as *M. aeruginosa* chlorophyll and photosynthetic quantum yield (p < 0.05) (Brown-Forsythe and Welch ANOVA assuming samples without equal variances). All macrophytes were compared to *P. crassipes* for biomass, whereas chlorophyll and yield were compared to the control condition. Dunnet’s post-hoc test and Tukey (biomass ratio) were applied to determine specific differences between treatments. The software used was Graph Pad Prism 8 version 8.0.2 (Boston, USA) for statistical analyses and chart plotting.

## 3. Results

### 3.1 Nutrient removal from the water by macrophytes

In the mesocosms, a significant reduction in dissolved nutrients (SRP, NH_4_^+^, and NO_3_^−^) was observed for all tested emergent macrophyte species compared to the control, where no plants were added to the water (Figure 2). *L. leptocarpa* presented the fastest reduction in SRP concentrations with values below the quantification limit (16 µg L^−1^) on the 7^th^ day. After 14 days, the SRP concentration in mesocosm water with *P. ferrugineum* and *U. mutica* was below the quantification limit, indicating a significant decrease compared to the control (p<0.05). *A. philoxeroides* and *T. domingensis* presented an SRP reduction of 94 % and 64 %, respectively, in 14 days (Figure 2A). In comparison, the floating macrophyte *P. crassipes*, used as a positive control for nutrient removal, reduced the SRP concentration by 86% after 14 days.

**Figure 2:**
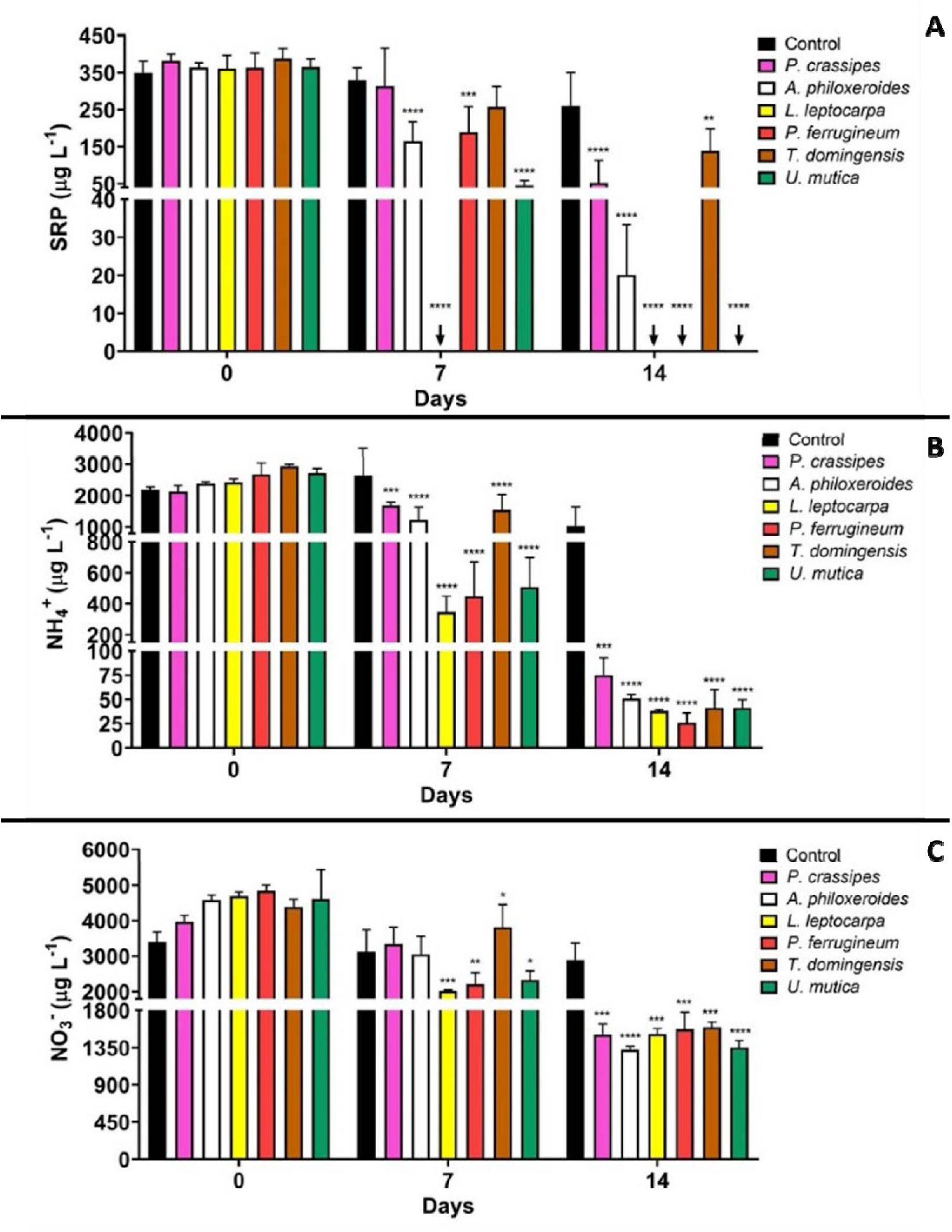
SRP, NH_4_^+^, and NO_3_^−^ concentrations in mesocosm water during macrophyte growth. Asterisks mean significant differences comparing each macrophyte to the control condition (mesocosm with no macrophyte) in the same sampling day, where * p<0.05, ** p<0.01, *** p<0.001, and *** p<0.0001. Arrows indicate values below the quantification limit (p<0.05).

The emergent macrophytes effectively removed NH_4_^+^ and NO_3_^−^ from the water on the 7^th^ day (except for *A. philoxeroides*) and intensified removal on the 14^th^ day (Figures 2B and C) (p<0.05). Compared to the control, in seven days *L. leptocarpa*, *P. ferrugineum*, and *U. mutica* showed the highest removal for NH_4_^+^ (86%, 83%, and 81%, respectively) and NO_3_^−^ (57 %, 54 %, and 49 %, respectively) (Figure 2B and C). Comparatively, the floating macrophyte *P. crassipes* removed 21 % and 16 % of NH_4_^+^ and NO_3_^−^in seven days.

TP concentrations in the water were significantly lower than in the control for all macrophytes (except *for T. domingensis* in 7 days). The strongest effect in TP decrease was observed for *L. leptocarpa* in 7 days (76 %) (Supplementary Figure 1A). No significant differences in TN concentrations were observed compared to the control, except for *T. domingensis* at day 14 (Supplementary Figure 1B). In mesocosms with the floating macrophyte *P. crassipes*, TP concentrations were lower than control on day 14, and TN was not different from control.

### 3.2 Water abiotic parameters

In all treatment groups (including *P. crassipes*), pH and DO values were significantly lower than control on days 7 and 14 (Figure 3). The initial pH value was 6.0, and all treatment groups decreased pH values to 4.0-5.0 after 14 days (Figure 3A). In the control condition, no variation was observed for DO, whereas, in the treatment groups, the DO values decreased until the 14^th^ day. The presence of *A. philoxeroides, P. ferrugineum*, and *U. mutica* resulted in the lowest DO values on the 14^th^ day (3.1 ± 0.2 mg L^−1^, 3.2 ± 0.2 mg L^−1^, and 1.8 ± 0.2 mg L^−1^, respectively) (Figure 3B). Water conductivity was not altered by the growth of macrophytes, except for *U. mutica* on the 14^th^ day (Supplementary Figure 2). During the experiment, the average water and air temperature values were 24.2 ± 0.3 °C and 27.0 ± 4.2 °C, respectively.

**Figure 3:**
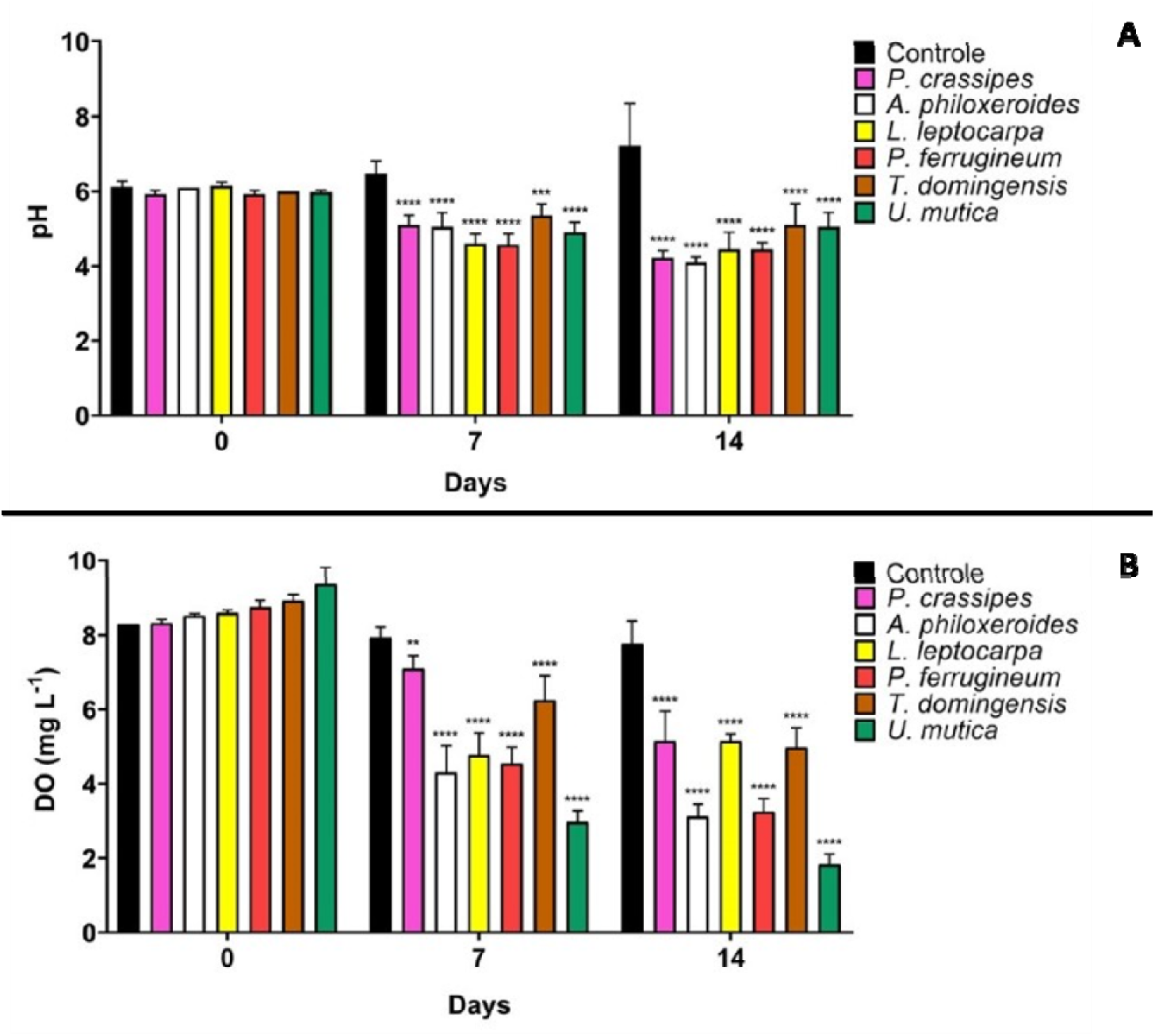
Variation of pH and DO concentration in mesocosm water during macrophyte growth. Asterisks mean significant differences comparing each macrophyte to the control condition (mesocosm with no macrophyte) on the same sampling day, where ** p<0.01, *** p<0.001, and **** p<0.0001.

### 3.3 Macrophyte and phytoplankton growth

During the experimental period, *P. crassipes* fresh biomass increased, reaching the highest ratio between final and initial biomass (59%) (Figure 4 and Supplementary Table 1). The emergent macrophytes increased biomass to a lesser extent after 14 days, achieving a percentage growth of: *A. philoxeroides* (45%), *L. leptocarpa* (24%), *P. ferrugineum* (24%), *U. mutica* (23%) and *T. domingensis* (16%) (Figure 4 and Supplementary Table 1).

**Figure 4:**
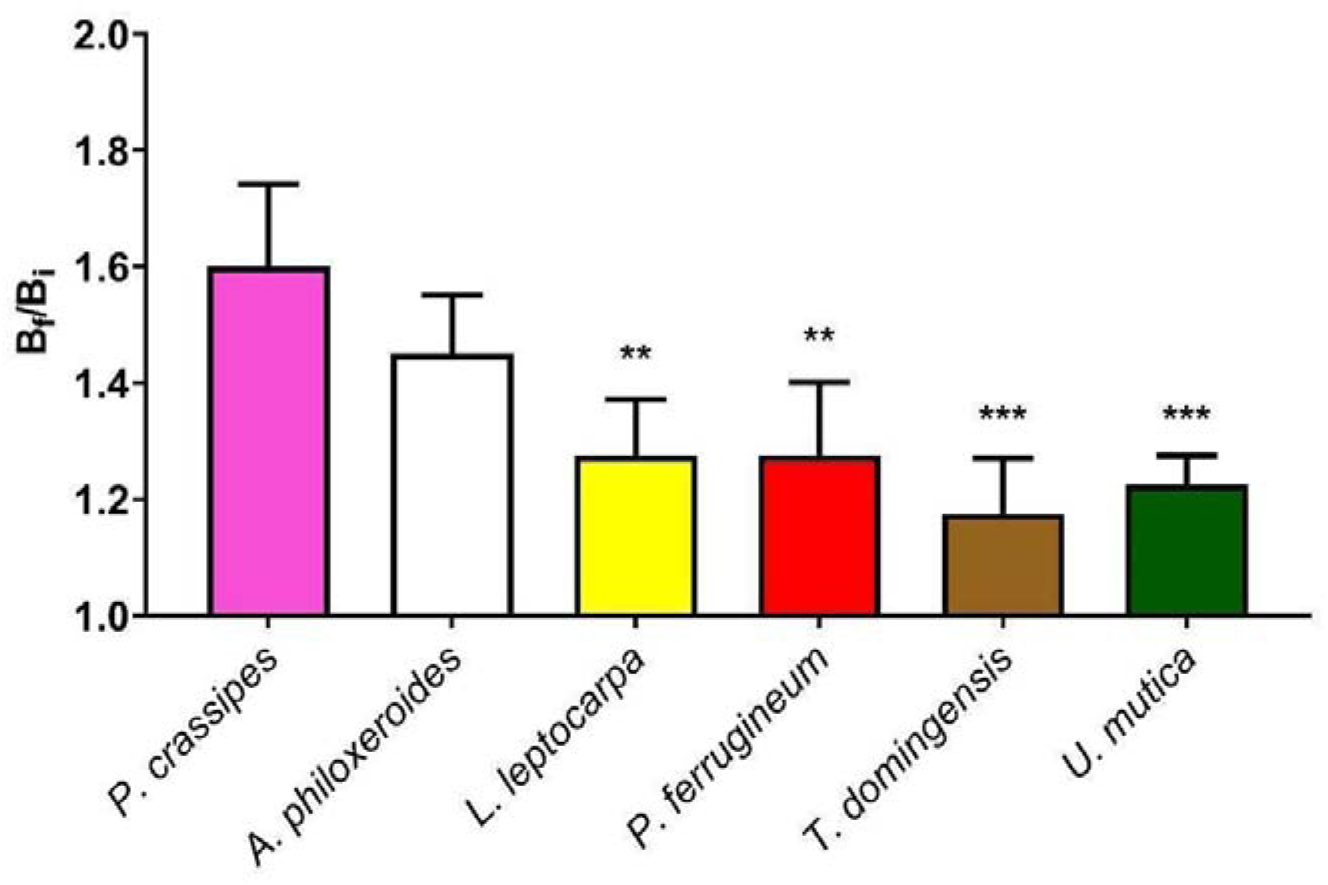
Ratio of macrophyte growth [Bf – final fresh biomass (g); Bi – initial fresh biomass (g)] after 14 days in the mesocosm system. Asterisks indicate a significant difference between all emergent macrophytes tested and *P. crassipes*, where \*\**p <* 0.01, \*\*\**p <* 0.001.

After seven days, the phytoplankton growth was similar in all treatment groups and in the control condition (Chl-a concentration of approximately 2.5 ± 1.2 µg L^−1^). On the 14^th^ day, Chl-*a* concentration increased to 30 µg L^−1^ in the control (Figure 5). In contrast, the mesocosms that received the emergent macrophytes and those with *P. crassipes* maintained the same Chl-*a* concentrations measured on the 7^th^ day (Figure 5, Supplementary Table 3).

**Figure 5:**
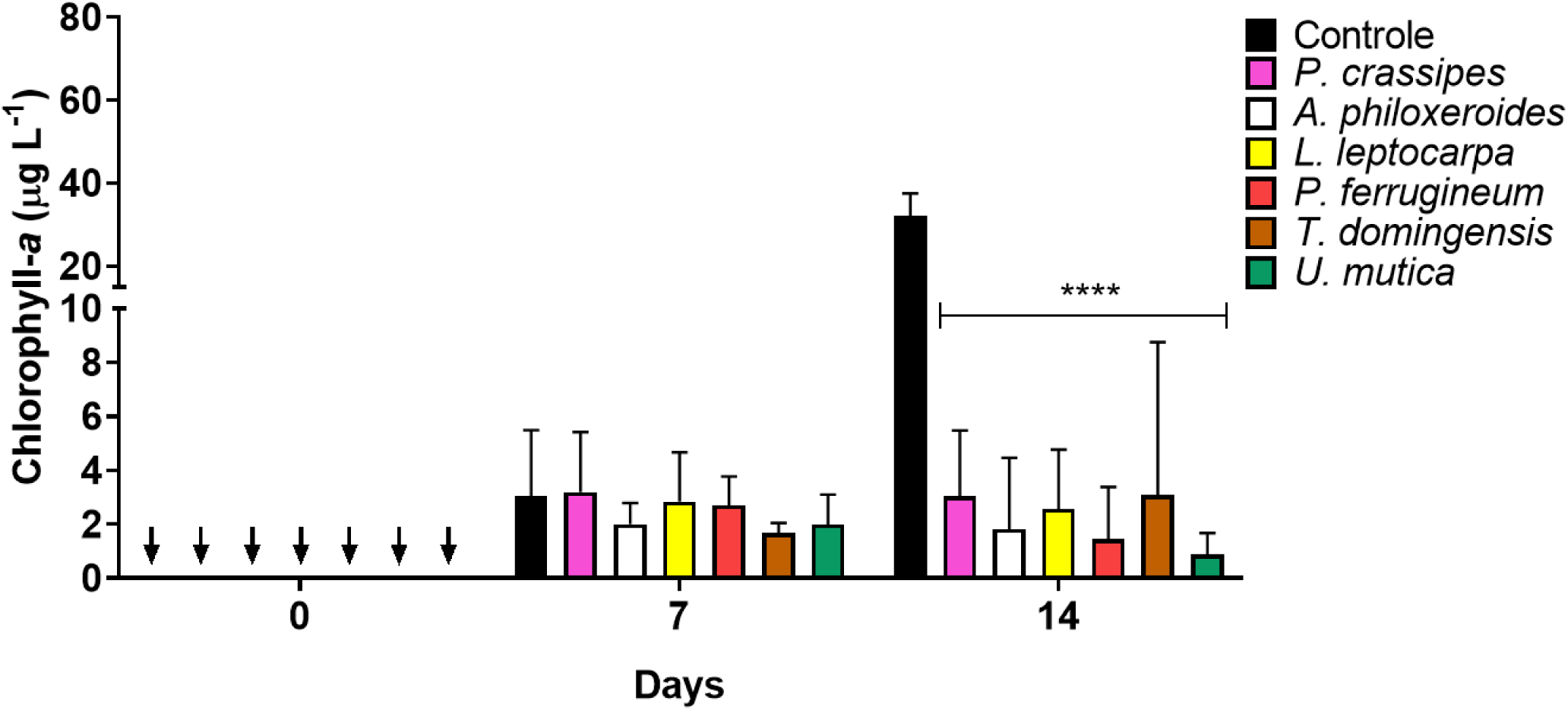
Variation of chlorophyll-*a* concentration in the mesocosm water during macrophyte growth. Asterisks significant differences comparing each macrophyte to the control condition (mesocosm with no macrophyte) in the same sampling day, where **** p<0.0001. Arrows indicate concentrations below the quantification limit (p<0.05).

The Chl-*a* concentrations were related to green algae, while cyanobacteria or diatoms were not detected.

### 3.4 Potential assimilation of dissolved phosphorus into macrophyte biomass

To estimate the potential assimilation of soluble reactive phosphorus (SRP) into the macrophyte biomass over 14 days, we related the decrease in SRP concentrations in water to the increase in plant biomass. These ratios resulted in similar values ranging from 2.7 to 3.2 without significant differences among the macrophyte species tested (p=0.85, F=0.38) (Figure 6).

**Figure 6:**
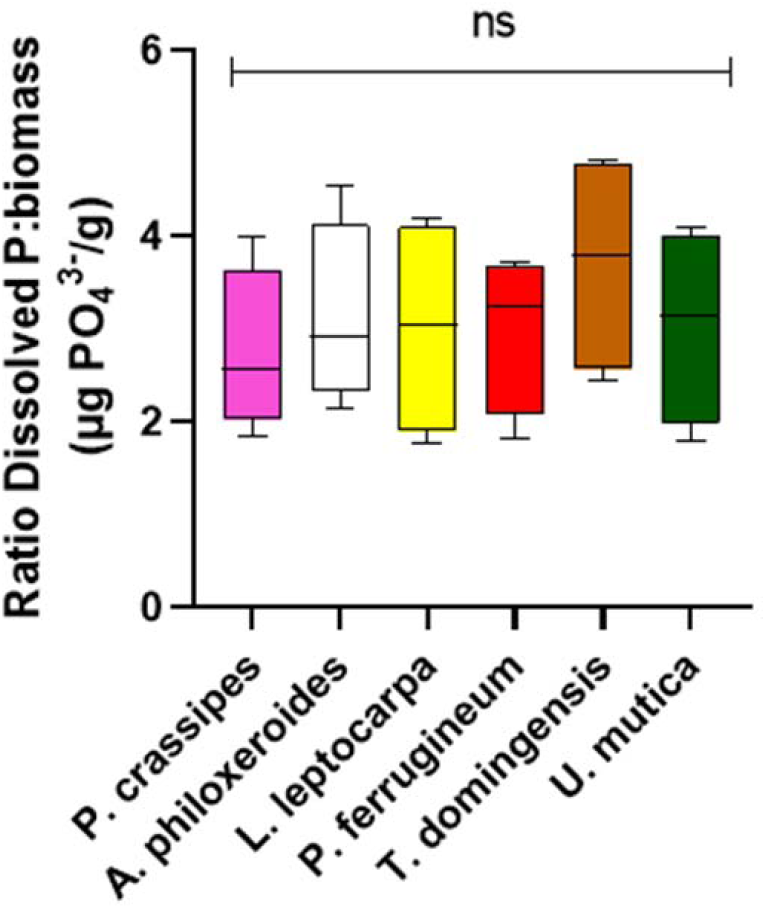
Ratio of dissolved phosphorus and macrophyte biomass (µg SRP/g biomass) over 14 days for each tested species, where ns means non-significant (p>0.05).

### 3.5 Allelopathic potential of root exudates against m. aeruginosa and characterization of their chemical profiles

Inhibition of *M. aeruginosa* growth was observed with the exudates of three macrophyte species: *A. philoxeroides, L. leptocarpa*, and *P. ferrugineum*, with Chl-*a* values below the detection limit on day 7 (Figure 7A). In the control condition (ASM-1) and incubation of *M. aeruginosa* with the exudates from *T. domingensis, U. mutica*, and *P. crassipes*, Chl-*a* concentrations increased approximately three times in this period compared with the initial concentration of chlorophyll, on average (dashed line) (Figure 7A). The efficiency of the photosynthetic activity was also suppressed by incubation with exudates from *A. philoxeroides, L. leptocarpa*, and *P. ferrugineum*, and only detected in the non-affected cultures (Figure 7B). Interestingly, the incubation of cyanobacterial cells with T. *domingensis* root exudate promoted a slight increase in the photosynthetic activity of *M. aeruginosa* compared to the control condition (p<0.05).

**Figure 7:**
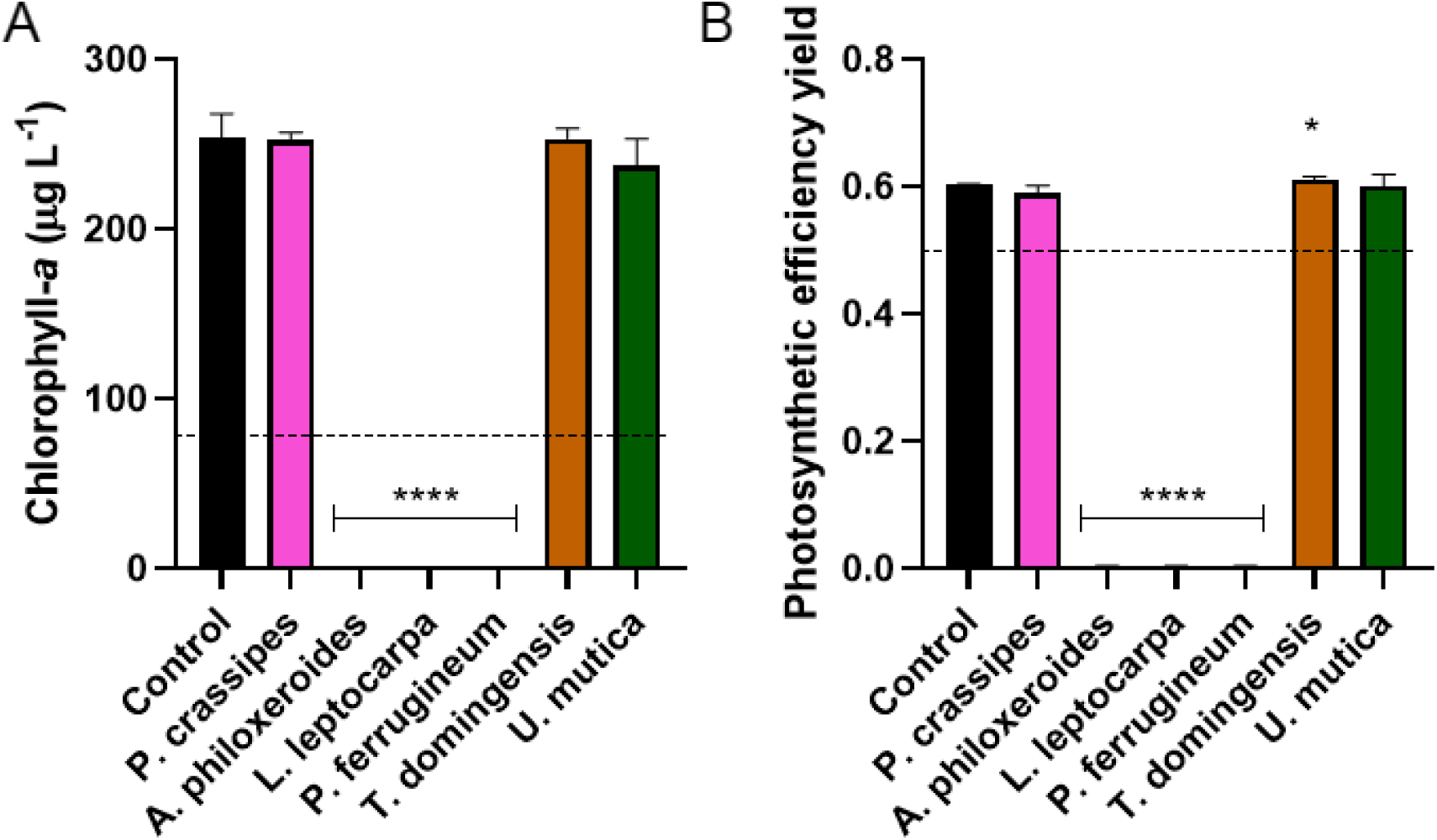
Estimated chlorophyll concentration (A) and photosynthetic activity (B) of *M. aeruginosa* cultures after 7 days of incubation with root exudates obtained from macrophytes. Dashed lines represent the average concentration of chlorophyll-*a* or yield at the initial time. *p<0.05 and **** p<0.0001.

The mass spectrometry analysis (FT-ICR/MS) was performed for those macrophytes that inhibited the growth of *M. aeruginosa*: *A. philoxeroides, L. leptocarpa,* and *P. ferrugineum*. Results indicated different molecular compositions of the exudates, as represented by the multivariate PDS-DA analysis considering both the positive and negative ionization modes (Supplementary Figure 3).

## 4. Discussion

Five emergent macrophytes tested for phytoremediation efficiency in mesocosm systems were adapted to experimental CFWs, and efficient to reduce P and N from the water column. It is interesting to note that the tested species represent a diversity of habitats, adaptive niches and taxonomic groups.

The shift in the cultivation of these plants, from emergent to floating in CFW, was not a limiting factor for their physiology as assessed by the removal of dissolved nutrients (SRP, NH_4_^+^, and NO_3_^−^), since the tested macrophytes were as efficient as the extensively used floating macrophyte *P. crassipes*. Among the emergent macrophytes, *L. leptocarpa* presented the fastest nutrient removal, reducing SRP concentrations to values below 16 µg L^−1^ on the 7^th^ experimental day, while *P. ferrugineum* and *U. mutica* reached this concentration on the 14^th^ day. The efficiency of NH_4_^+^ and NO_3_^−^ removal by *L. leptocarpa* was similar to *P. ferrugineum* and *U. mutica* in the same period.

It should be noted that N and P removal is also attributed to the symbiotic relationship with root microbes, constituting ‘the macrophyte-rhizosphere complex’ which is essential for this ecosystem service (Rejmankova, 2011; Rheman et al., 2017). Although we estimated SRP removal as P assimilation, it would be necessary to consider the microbial uptake associated with the ectorhizosphere. The possible diversity and differential functionality of the macrophyte-rhizosphere complex of the different species should be considered in future investigations. Additionally, apart from active assimilation, the physical adsorption of P to the roots by interaction with charged particles can participate in the decrease of P concentrations in the water (Devau et al., 2010).

Except for *A. philoxeroides*, all tested macrophytes showed lower growth than *P. crassipes* during the 14-day incubation, indicating a high nutrient removal ability of emergent macrophytes in CFWs. *T. domingensis* was the macrophyte with the lowest growth in mesocosms, yet it was equally efficient to all tested species in removing NH_4_^+^ and NO_3_^−^ (Figure 2). Removal of SRP by this species was slower, suggesting that the macrophyte requires more P in a later growth stage. Interestingly, all macrophytes presented a similar SRP: biomass ratio over 14 days of incubation. This observation suggests that this period was enough to stabilize their biomass considering the initial availability of dissolved phosphorus. However, in *P. crassipes*, *A. philoxeroides,* and *T. domingiesis* mesocosms, SRP concentrations were still detected in the water after 14 days (~50, 20 and 150 µg L^−1^, respectively). Thus, smaller macrophytes that absorb dissolved P faster, such as *L*. *leptocarpa*, *P. ferrugineum,* and *U. mutica*, would be more beneficial for P phytoremediation in CFWs.

Considering dissolved nitrogen forms, NH_4_^+^ concentrations had a higher reduction than NO_3_^−^ concentrations. This can be explained by the dynamics of N absorption in the roots. NO_3_^−^ is actively absorbed by roots with ATP consumption, while NH_4_^+^ is absorbed by facilitated transport without energy consumption (Reid and Hayes, 2003). In addition, NO_3_^−^ was detected in a higher initial concentration in the mesocosm water than NH_4_^+^, and the removal could be affected by the saturation of N in macrophyte tissues, as indicated by NO_3_^−^ concentrations in mesocosms with *L. leptocarpa*, *P. ferrugineum* and *U. mutica*, where the removal of NO_3_^−^ was faster until the 7^th^ day and decreased from 7 to 14 days.

In the mesocosms maintained with macrophytes, phytoplankton growth was inhibited. The primary effect could be associated with SRP availability that decreased in mesocosms containing *L. leptocarpa, P. ferrugineum*, and *U. mutica* reaching limiting concentrations for phytoplankton growth (lower than the detection limit of 16 µg L^−1^) (Lewis and Roberson, 2022). In all treatment groups, dissolved N (NH_4_^+^ + NO_3_^−^) concentrations did not reach limiting concentrations for phytoplankton (20 µg L^−1^) (Reynolds, 2006). Other studies showed that CFW systems efficiently controlled phytoplankton growth by nutrient limitation (West et al., 2017; Benvenuti et al., 2018). Additionally, it is important to point out that light was not limited to phytoplankton, as the platforms occupied half of the mesocosm surface. This was evident in the control condition containing only platforms without plants, where phytoplankton growth was observed over time.

The excessive biomass production of the floating macrophyte *P. crassipes* is a challenge for its management (Patel, 2012; Djihouessi et al., 2023). In this study, *P. crassipes* had the highest biomass gain in 14 days (59%). In contrast, except for *A. philoxeroides*, the emerging species tested in this study increased their biomass by less than 25%. Even producing less biomass in this period, *L. leptocarpa*, *P. ferrugineum*, and *U. mutica* removed dissolved nutrients (SRP, NH_4_^+^, and NO_3_^−^) in a shorter time and/or higher amount compared to *P. crassipes*. Moreover, more controlled growth means easier management, which is a positive feature for using these emergent macrophytes in CFW.

The observed allelopathic effect caused by the root exudate of *A. philoxeroides, L. leptocarpa*, and *P. ferrugineum* on *M. aeruginosa* is an attractive characteristic and increases the repertoire of macrophytes that present allelopathic effects on cyanobacteria (Mohamed, 2017). Our findings corroborate the studies carried out by Zuo et al. (2011 and 2012), which reported that green algae (the Chlorophyceae *Chlorella pyrenoidosa*) and cyanobacteria (*Microcystis aeruginosa*) were inhibited by exudates and extracts of *A. phyloxeroides* under both laboratory and in-situ conditions. For *L. leptocarpa* and *P. ferrugineum,* as far as we know, this is the first report documenting the inhibitory effects of these macrophytes on a cyanobacterial species. Studies on larger scales with natural cyanobacteria blooms can help better understand the extent of these observed effects in vitro.

The FT-ICR-MS multivariate analysis (PLS-DA) suggested a different molecular composition for the root exudates that inhibited *M. aeruginosa* growth. This is an expected result since these macrophytes have a great phylogenetic distance (Table 1). Probably each macrophyte species presents a specific molecular repertoire, which would result in the inhibition of *M. aeruginosa* growth. Although allelopathic activity on cyanobacteria was described for several species of macrophytes (emergent, floating and submerged) the underlying mechanisms of inhibition were characterized for only a fraction of these (Mohamed, 2017). The investigation of a larger number of macrophytes may reveal new forms of inhibition of cyanobacterial physiology.

In addition to nutrient removal and allelopathic effects, CFWs cause changes in abiotic parameters such as water pH which may affect the phytoplankton. The acidification observed in the mesocosm water was caused by the release of CO_2_ from the macrophyte roots and its associated microbiota (Hanson et al., 2000), a product of their heterotrophic activity. A pH reduction (below 7.0) limits the growth of cyanobacteria, as they require the inorganic form of carbon HCO ^−^, which is formed in alkaline pH, reaching its maximum saturation in water between a pH range 8-9 (Mangan et al., 2016). However, the extent of acidification caused by the macrophytes and its potential detrimental effect on cyanobacteria could be better explored in full-scale applications.

One of the primary purposes of CFW systems should be to limit P availability in aquatic environments. In freshwater, P availability is considered a limiting factor for phytoplankton growth, mainly in tropical environments, compared to N-rich compounds (Huszar et al., 2006). Furthermore, different from N, which is available in the atmosphere and can be fixed by microbial metabolism, P is obtained from mineral sources and is becoming scarce for agriculture worldwide. A significant part of P is lost in the aquatic environment, causing eutrophication, and is not recycled (Cooper et al., 2011). CFW systems could be an efficient nature-based solution to recover P from aquatic environments and return it as a biofertilizer for crops or reforestation, representing a valuable tool in the circular economy (Quilliam et al., 2015; Poveda, 2022; Waylen et al., 2024). Considering the high nutrient absorption activity of native macrophytes and their allelopathic effects, CFWs could be interesting tools to deliver ecosystem services of eutrophication control and suppression of phytoplankton blooms.

## 5. Conclusion

Five emergent macrophytes with wide distribution in the American continent were adapted to CFW. They efficiently removed nutrients from the water column in a way equivalent or superior to the commonly used floating macrophyte *P. crassipes*. Additionally, root exudates from *A. philoxeroides*, *L. leptocarpa*, and *P. ferrugineum* presented allelopathic activity on *M. aeruginosa,* which was not observed for *P. crassipes*. Using these emergent species in CFW systems is a viable alternative approach to controlling eutrophication in natural environments and potentially controlling cyanobacteria blooms.

## Acknowledgments

We acknowledge all funding support from CNPQ (proc. number <u>307219/2022-4</u>), FAPERJ (proc. number E-26/200.551/2023 (281293)) and ELERA Renováveis SA (contract number 7255) as well as Dra. Evelyn Maribel Condori Peñaloza for helping us with FT-ICR-MS analysis and interpretation.

## Data availability statement

All data will be available on request from the authors. The data that support the findings of this study are available from the corresponding author, C.M.L.F upon reasonable request.

## Supplementary materials

Characterization of macrophyte root exudates by high-resolution mass spectrometry

Experiments were carried out on a Bruker SolariX XR, 7 Tesla Fourier Transform Ion Cyclotron Resonance Mass Spectrometry (FT-ICR-MS) (Brüker Daltonics Inc., Billerica, MA) equipped with an electrospray ionization (ESI) source. The samples and blanks were directly infused at a flow rate of 3.0 μL min^−1^. The ESI source parameters in positive and negative ionization modes were nebulizer gas pressure of 1.0 bar, dry temperature of 200 °C, dry gas of 4 L min^−1,^ and capillary potential of 5.0 kV. The spectra were acquired with a mass range from 75.26 to 1200 (m/z) (with a resolution of 200 000 at m/z 400), 10 scans accumulated for each sample, and mass accuracy of < 2 ppm, provided the unambiguous molecular formula assignments for singly charged molecular ions. Elemental compositions of the compounds were designed by measuring the m/z value. The unsaturation level of each molecule was deduced directly from its double bond equivalent (DBE), following the equation DBE = c - h/2 + n/2 + p/2 + 1, where c, h, n, and p are the letters of carbon, hydrogen, nitrogen, and phosphorous atoms, respectively.

Data were normalized using the MetaboAnalyst 4.0 platform (Chong et al., 2019). Multivariable analysis partial least squares-discriminant analysis (PLS-DA) was performed with the MetaboAnalyst 4.0 platform to distinguish the root exudates of the species that presented allelopathic activity.

**Supplementary Table 1:**
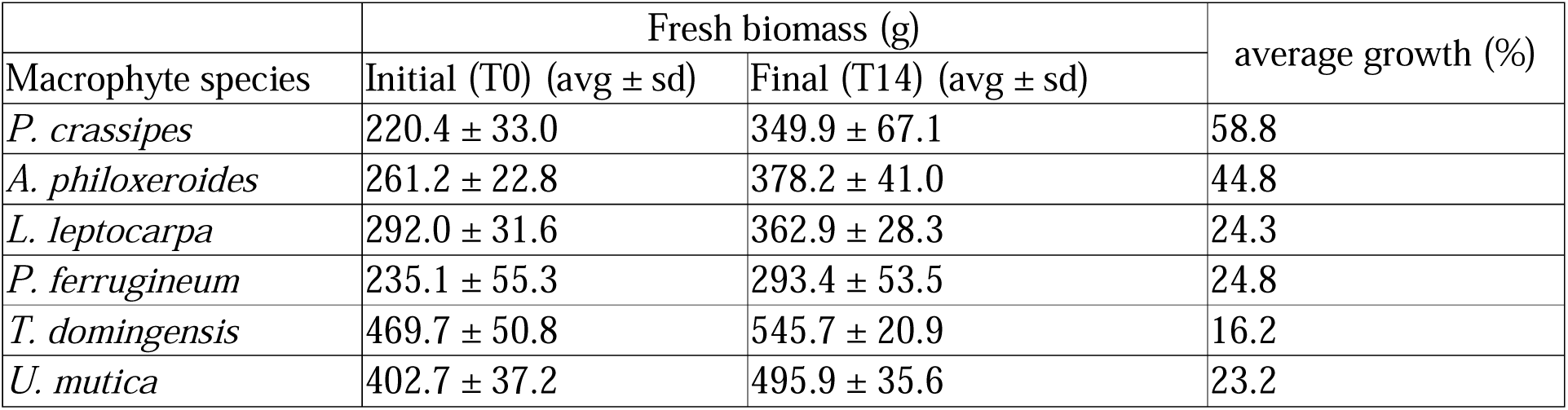
Variation of fresh biomass and percentage growth (average) of macrophytes over 14 days in mesocosms.

**Supplementary Figure 1:**
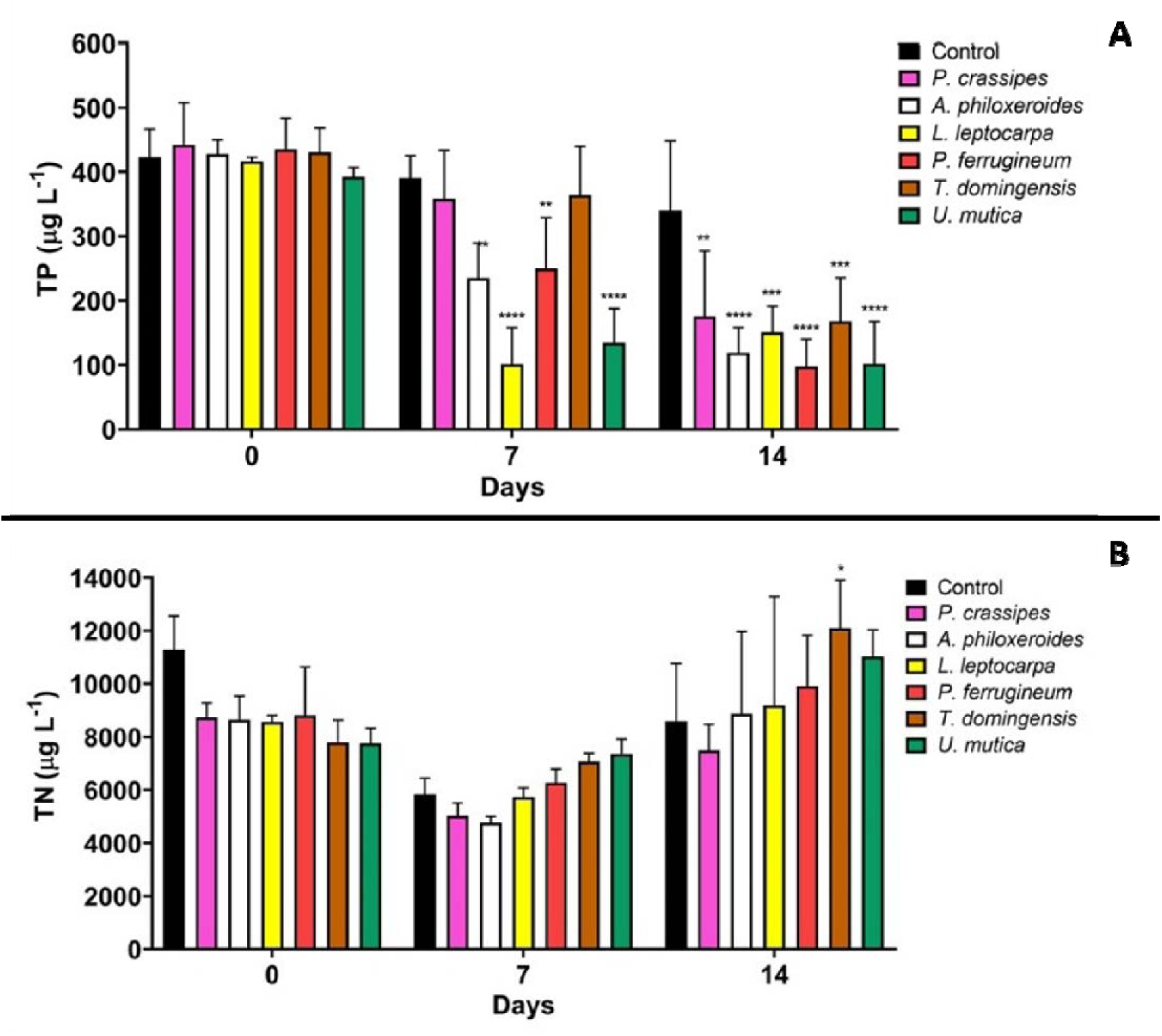
TP and TN concentrations in mesocosm water during macrophyte growth. (p<0.05).

**Supplementary Figure 2:**
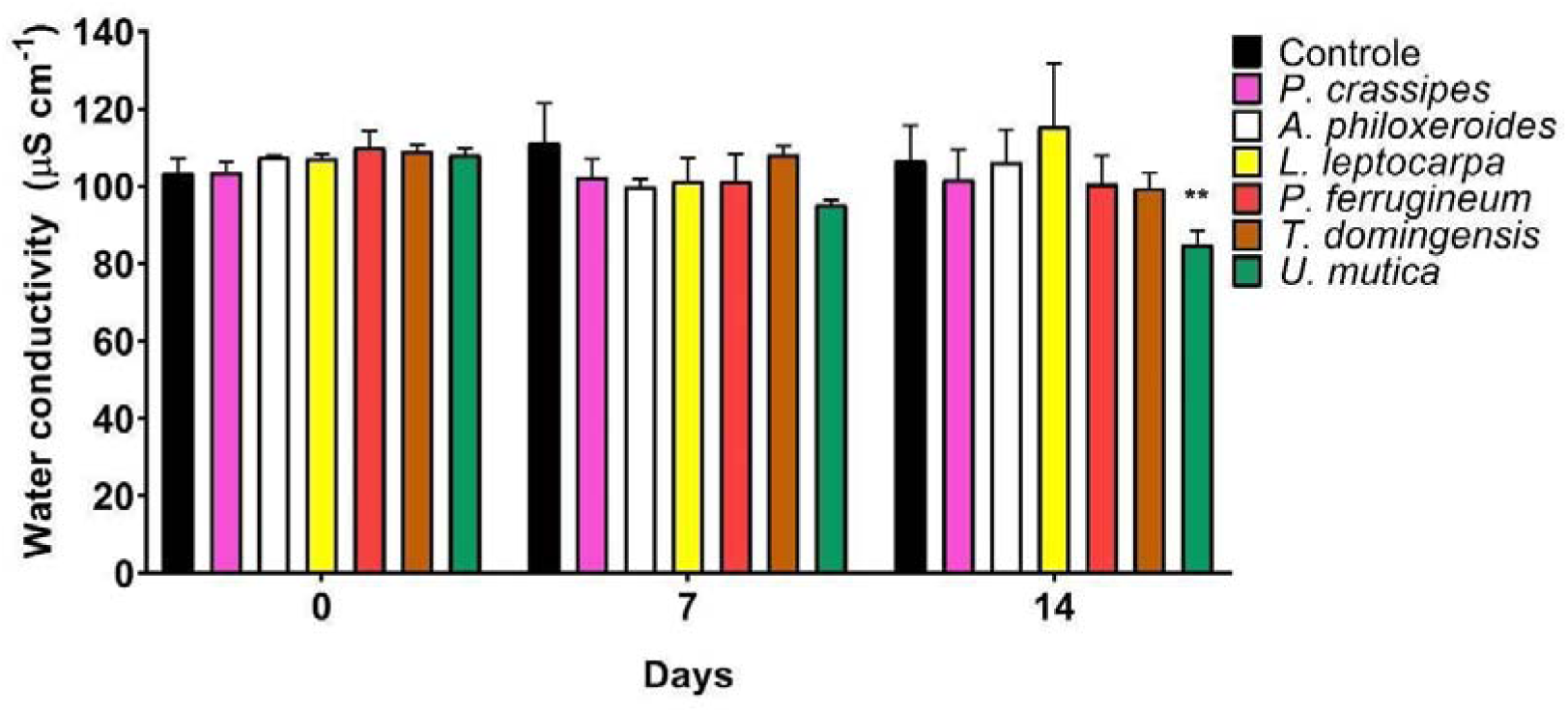
Conductivity in mesocosm water during macrophyte growth (p<0.05).

**Supplementary Figure 3:**
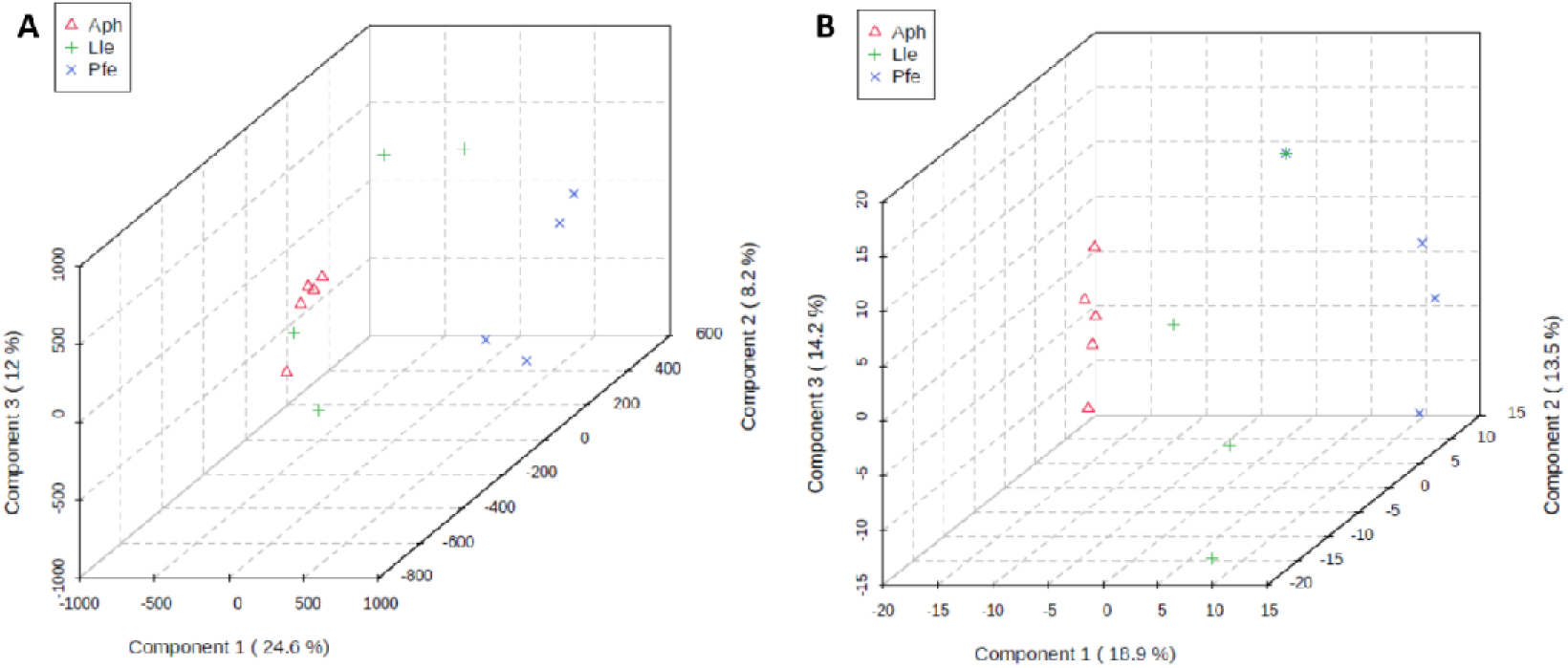
Multivariate analysis (PLS-DA) of the compounds identified in root exudates of *A. philoxeroides* (Aph), *L. leptocarpa* (Lle) and *P. ferrugineum* (Pfe) using mass spectrometry. (A) Positive mode, (B) Negative mode.

