## Supplementary Materials for "WHO WINS? COMPARISON BETWEEN EMERGENT MACROPHYTES AND PONTEDERIA CRASSIPES (WATER HYACINTH) AS NATURE-BASED SOLUTIONS FOR NUTRIENT PHYTOREMEDIATION AND CYANOBACTERIA MITIGATION"

Table S1: Percentual removal of nutrients or inhibition of phytoplankton (chlorophyll content) by all macrophytes related to the control condition respectively to time 7 and 14 days. The dashes mean any removal or inhibition could be calculated. Results are shown in average ± standard deviation (%).

|  | PO_4_^-3^ | | P total | | NH_4_^+^ | | NO_3_^-^ | | N total | | Chl | |
| --- | --- | --- | --- | --- | --- | --- | --- | --- | --- | --- | --- | --- |
|  | 7 d | 14 d | 7 d | 14 d | 7 d | 14 d | 7 d | 14 d | 7 d | 14 d | 7 d | 14 d |
| ***Pontederia crassipes*** | 3±34 | 76±31 | 8±21 | 41±40 | 30±26 | 90±5 | - | 47±13 | 14±8 | 15±28 | - | 92±3 |
| ***Alternanthera philoxeroides*** | 48±17 | 91±8 | 39±19 | 62±17 | 45±35 | 93±4 | - | 53±6 | 17±11 | 4±27 | - | 89±18 |
| ***Ludwigia leptocarpa*** | 97±5 | 96±4 | 74±15 | 52±20 | 84±12 | 95±3 | 34±12 | 47±7 | - | - | - | 93±3 |
| ***Polygonum ferrugineum*** | 41±28 | 99±2 | 34±26 | 72±8 | 83±6 | 97±2 | 27±20 | 45±3 | - | - | - | 94±8 |
| ***Typha domingensis*** | 20±22 | 40±37 | 6±19 | 42±39 | 13±13 | 94±6 | - | 44±11 | - | - | 11±69 | 96±8 |
| ***Urochloa mutica*** | 87±3 | 98±3 | 66±11 | 72±12 | 76±21 | 95±3 | 23±22 | 52±10 | - | - | - | 96±6 |

Table S2: Variation of fresh biomass and percentage growth (average) of macrophytes over 14 days in mesocosms.

|  | Fresh biomass (g) | | average growth (%) |
| --- | --- | --- | --- |
| Macrophyte species | Initial (T0) (avg ± sd) | Final (T14) (avg ± sd) |  |
| *P. crassipes* | 220.4 ± 33.0 | 349.9 ± 67.1 | 58.8 |
| *A. philoxeroides* | 261.2 ± 22.8 | 378.2 ± 41.0 | 44.8 |
| *L. leptocarpa* | 292.0 ± 31.6 | 362.9 ± 28.3 | 24.3 |
| *P. ferrugineum* | 235.1 ± 55.3 | 293.4 ± 53.5 | 24.8 |
| *T. domingensis* | 469.7 ± 50.8 | 545.7 ± 20.9 | 16.2 |
| *U. mutica* | 402.7 ± 37.2 | 495.9 ± 35.6 | 23.2 |

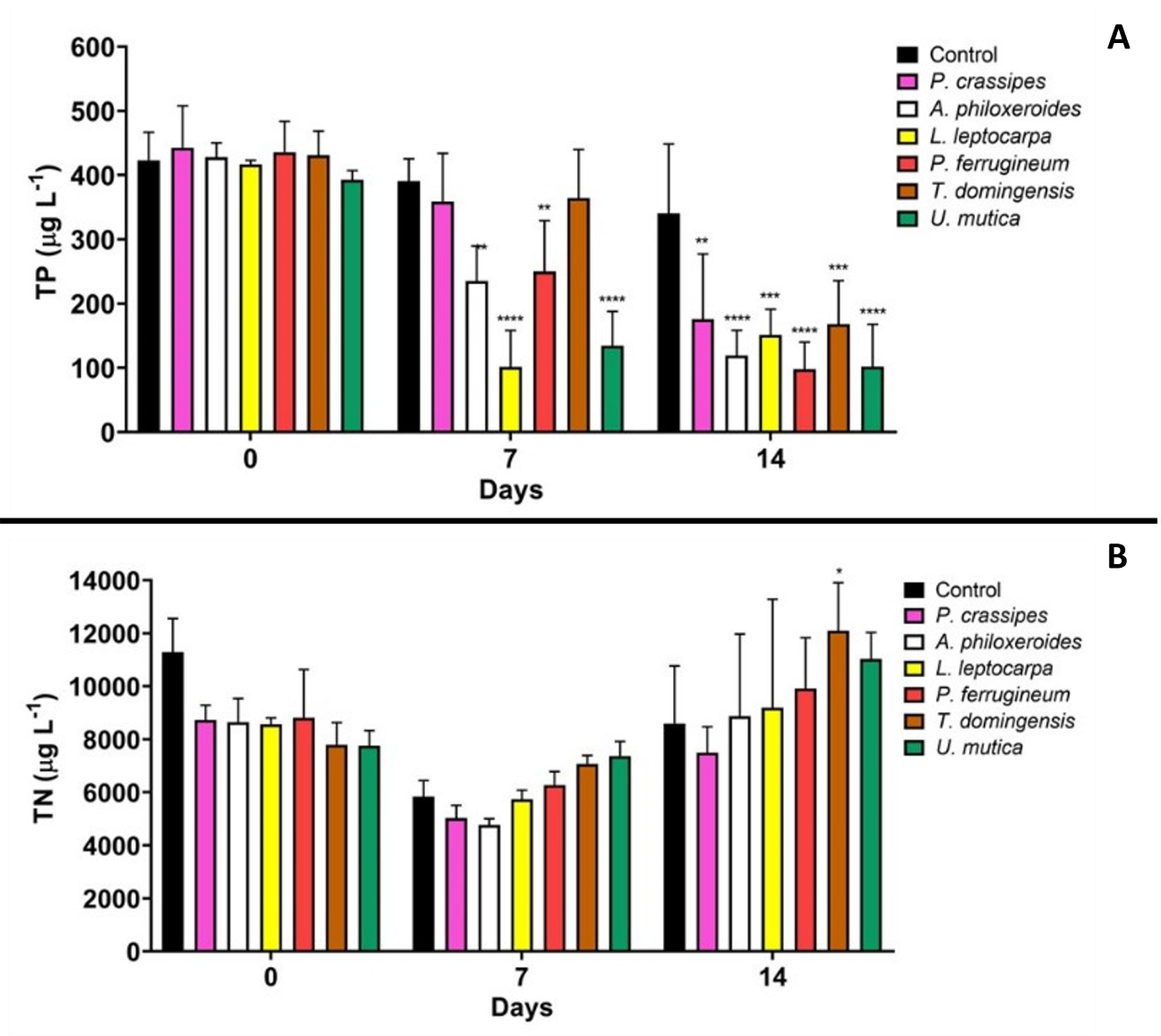
Figure S1: TP (A) and TN (B) concentrations in mesocosm water during macrophyte growth. (p<0.05).

.

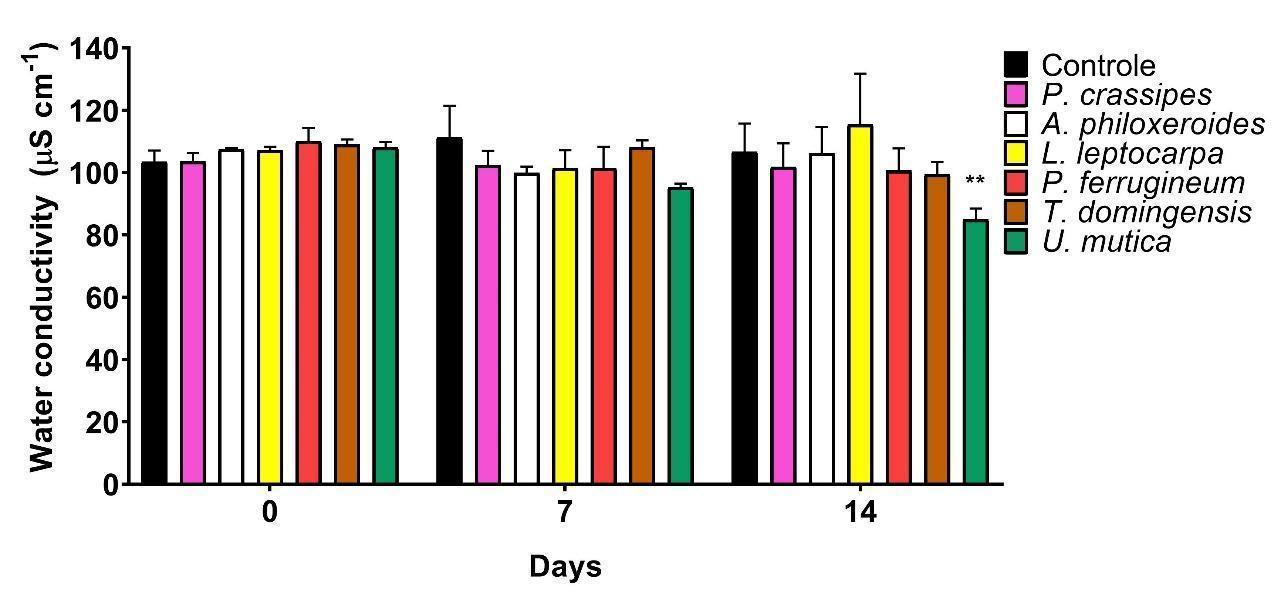
 Figure S2: Conductivity in mesocosm water during macrophyte growth (p<0.05).

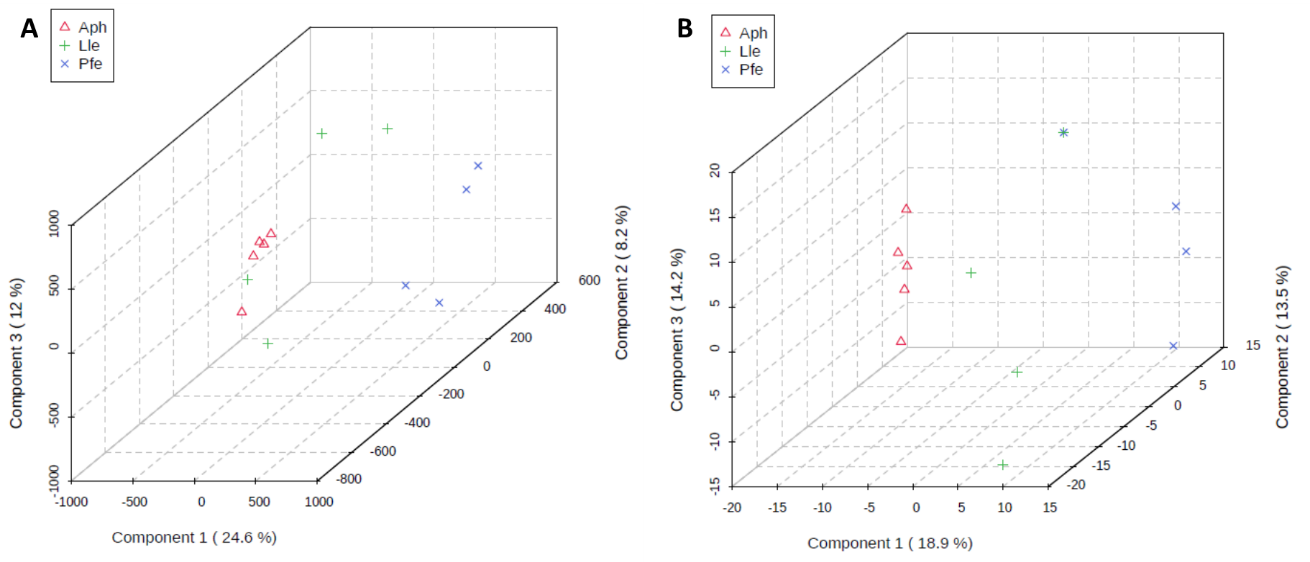

Figure S3: Multivariate analysis (PLS-DA) of the compounds identified in root exudates of *A. philoxeroides* (Aph), *L. leptocarpa* (Lle) and *P. ferrugineum* (Pfe) using mass spectrometry. (A) Positive mode, (B) Negative mode.
